# The Crown Pearl V2: an improved genome assembly of the European freshwater pearl mussel *Margaritifera margaritifera* (Linnaeus, 1758)

**DOI:** 10.1101/2023.02.11.528107

**Authors:** André Gomes-dos-Santos, Manuel Lopes-Lima, André M. Machado, Thomas Forest, Guillaume Achaz, Amílcar Teixeira, Vincent Prié, L. Filipe C. Castro, Elsa Froufe

**Author notes:** Corresponding authors: André Manuel Gomes dos Santos; Elsa Froufe.

## Abstract

Contiguous assemblies are fundamental to decipher the exact composition of extant genomes. In molluscs, this task is considerably challenging owing to their large size, heterozygosity, and widespread content of repetitive content. Consequently, the usage of long-read sequencing technologies is fundamental to achieve high contiguity and quality. The first genome assembly of *Margaritifera margaritifera* (Linnaeus, 1758) (Mollusca: Bivalvia: Unionida), a culturally relevant, widespread, and highly threatened species of freshwater mussels, has been produced recently. However, the current genome is highly fragmented since the assembly relied solely on short-read approaches. To overcome this caveat, here, a new improved reference genome assembly is produced using a combination of PacBio CLR long reads and Illumina paired-end short reads. This novel genome assembly is 2.4 Gb long, organized into 1,700 scaffolds with a contig N50 length of 3.4Mbp. The *ab initio* gene prediction resulted in a total of 48,314 protein-coding genes. This new assembly represents a substantial improvement and is an essential resource for studying this species’ unique biological and evolutionary features that ultimately will help to promote its conservation.

## Background and context

Initial efforts to sequence molluscan genomes relied primarily on short-read approaches, which, despite their unarguable value, frequently result in highly fragmented assemblies [1–4]. Consequently, long-read sequencing approaches, such as Pacific Bioscience (PacBio) or Nanopore (Oxford Nanopore), are becoming the common ground of emerging molluscan genome assembly projects [1–4]. This is further facilitated by the decreasing cost trend, coupled with increasing sequencing accuracy of these approaches [5]. Additionally, the structural information provided by long-reads is crucial to span large indels or inform about long structural variants (e.g., [6–8]), which is particularly relevant for molluscans that have large, heterozygous and highly repetitive genomes (reviewed in [4]). Consequently, long-read-based reference assemblies have reduced levels of fragmentation, fewer levels of missing and truncated genes, and reduced chances of chimerically assembled regions [6,7].

Bivalves from the Order Unionida, commonly known as freshwater mussels, are the most diverse group of strictly freshwater bivalves, with over 1,000 species distributed across all continents, except Antarctica [9,10]. The freshwater pearl mussel, *Margaritifera margaritifera* (Linnaeus, 1758) is perhaps the most emblematic, culturally significant, and known species of freshwater mussel. The freshwater pearl mussel is also the only species of the group that inhabits both European and North American freshwater systems [11,12] (Figure 1), mainly in cool oligotrophic waters. Moreover, this species holds a series of distinctive biological features, such as the ability to produce pearls (with an ancient history of pearl harvesting [13,14]), a long lifespan (reaching over 200 years [15]) with negligible signs of cellular senescence [16], and, as all other freshwater mussels, an obligatory parasitic life stage on salmonid fish species [12,17]. In the past, the freshwater pearl mussel was highly abundant across its Holarctic distribution [12]. However, during the last century, the species has suffered massive declines as a result of the many human-mediated threats that impact the freshwater ecosystems [11,12]. As a result, the species is listed as critically endangered in Europe and included in the European Habitats Directive under Annexes II and V, and the Appendix III of the Bern Convention [11].

**Figure 1.**
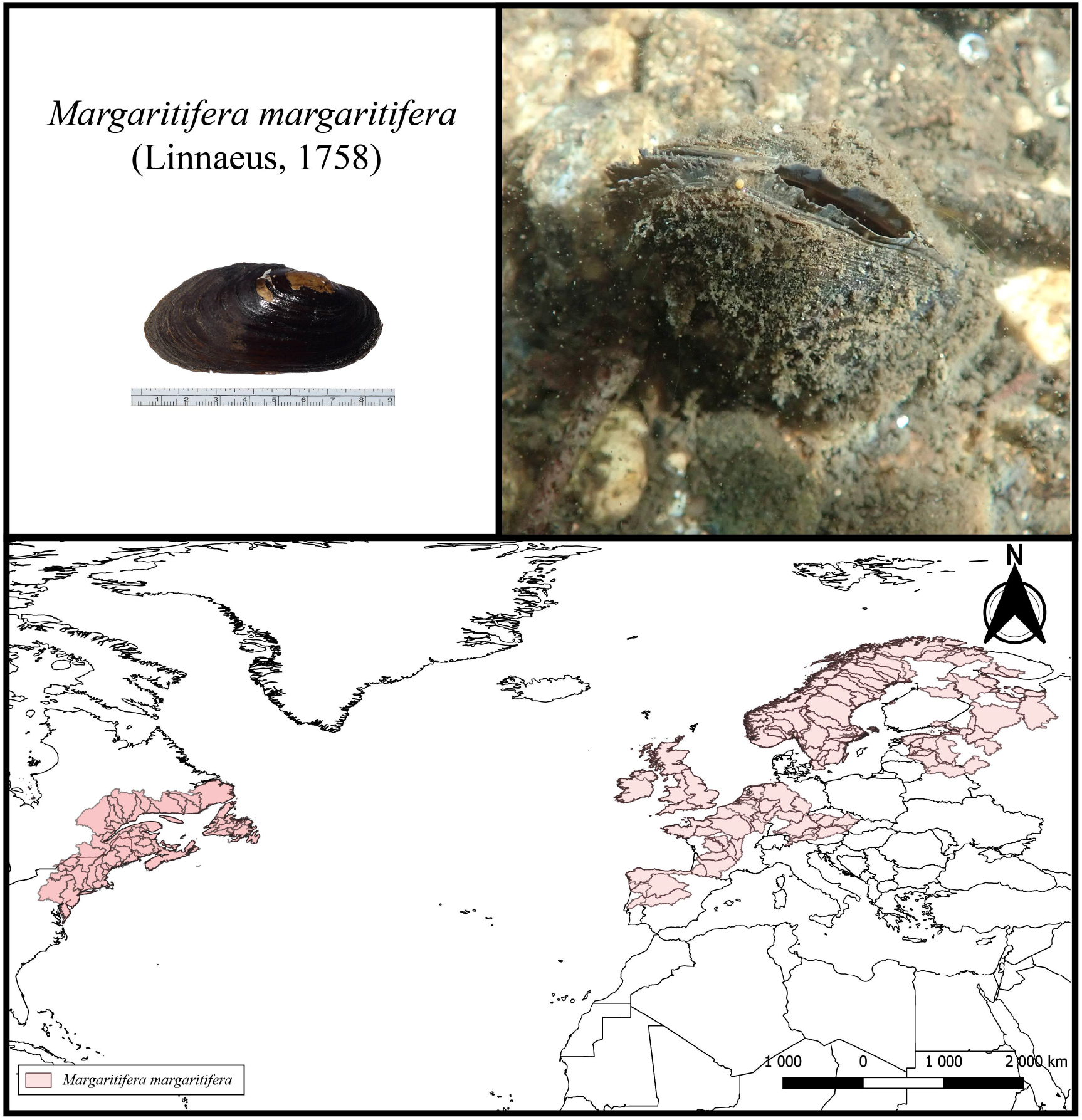
Top left: The *Margaritifera margaritifera* specimen used for the whole genome assembly. Top right: A specimen of *Margaritifera margaritifera* in its natural habitat (Photos by André Gomes-dos-Santos). Bottom: Map of the potential distribution of the freshwater pearl mussel, produced by overlapping points of recent presence records (obtained from [11]) with Hydrobasins level 5 polygons [26]. The potential distribution for Europe was retrieved from [11] and for North America from [27].

Despite the cultural significance and poor conservation status of the freshwater pearl mussel, the availability of genomic resources to study this species is still limited [13,18–22] and almost nothing is known about the molecular mechanism that governs the regulation and functioning of its many relevant biological features. Genomic resources provide benchmarking tools to monitor, identify, and classify conservation units as well as classify genetic elements with conservation relevance and adaptive potential [23,24], thus representing invaluable tools to improve the success of conservation efforts. Consequently, the sequencing of the first genome for the freshwater pearl mussel represented a fundamental resource for the study of its biology and evolution and, ultimately, promote its conservation [13]. Although the quality of this first assembly is good (validated with several statistics), the fact that it was produced solely using short-read sequencing (i.e., Illumina paired-end and mate-pair sequencing), hampered its overall contiguity [13]. The subsequent release of the highly contiguous genome assembly of the freshwater mussel *Potamilus streckersoni* (Smith, Johnson, Inoue, Doyle & Randklev, 2019), which relied on PacBio long-read sequencing, demonstrated how the use of longer reads is critical to ensure improved contiguity of genome assemblies for the group [25].

Aiming to improve the genome assembly for the freshwater pearl mussel, *M. margaritifera*, here, the genome of a new individual is sequenced using PacBio CLR and Illumina paired-end short reads. This new assembly represents the most contiguous freshwater mussel genome assembly available to date, representing a significant improvement in the contiguity and completeness [13].

## Methods

### Animal sampling

One individual of *M. margaritifera* was collected from the Tuela River in Portugal (Table 1) and transported alive to the laboratory, where tissues were separated, flash-frozen, and stored at −80□°C. The shell and tissues are deposited at CIIMAR tissue and mussels’ collection.

**Table 1.** MixS Descriptor for the freshwater pearl mussel *Margaritifera margaritifera* specimen used for the whole genome sequencing.

| Sample | *Margaritifera margaritifera* |
| --- | --- |
| Investigation_type | Eukaryote |
| Lat_lon | 41.862414; -6.931596 |
| Geo_loc_name | Portugal |
| Collection_date | 06/07/2021 |
| Env_package | Water |
| Collector | Amílcar Teixeira |
| Sex | Undetermined |
| Maturity | Mature |

### DNA extraction and sequencing

For PacBio sequencing, mantle tissue was sent to Brigham Young University (BYU, USA), where high-molecular-weight DNA extraction was performed and PacBio library construction was achieved following the SMRT bell construction protocol. The library was sequenced on a single-molecule real-time (SMRT) cell of a PacBio Sequel II system v.9.0. Genomic DNA for short-read sequencing was extracted from muscle tissue using the Qiagen MagAttract HMW DNA Kit, following the manufacturer’s instructions. The extracted DNA was sent to Macrogen Inc. for standard Illumina Truseq Nano DNA library preparation and whole genome sequencing of 150□bp paired-end reads on the Illumina Novaseq 6000 machine (Table 2).

**Table 2.** General statistics of raw sequencing reads used for the *Margaritifera margaritifera* genome assembly.

| Sample | Sequencing type | Library type | Platform | Insert size (bp) | Number of reads | Application |
| --- | --- | --- | --- | --- | --- | --- |
| PacBio CLR | WGS | Long Reads | PacBio Sequel II System | 14,339 | 7,892,056 | Genome Assembly, Genome Polishing |
| Illumina PE | WGS | Short Reads | Novaseq 6000 | 450 | 1,369,564,530 | Genome size estimation, Genome Polishing |

### Genome assembly and annotation

The overall pipeline used to obtain the genome assembly and annotation is provided in Figure 2.

**Figure 2.**
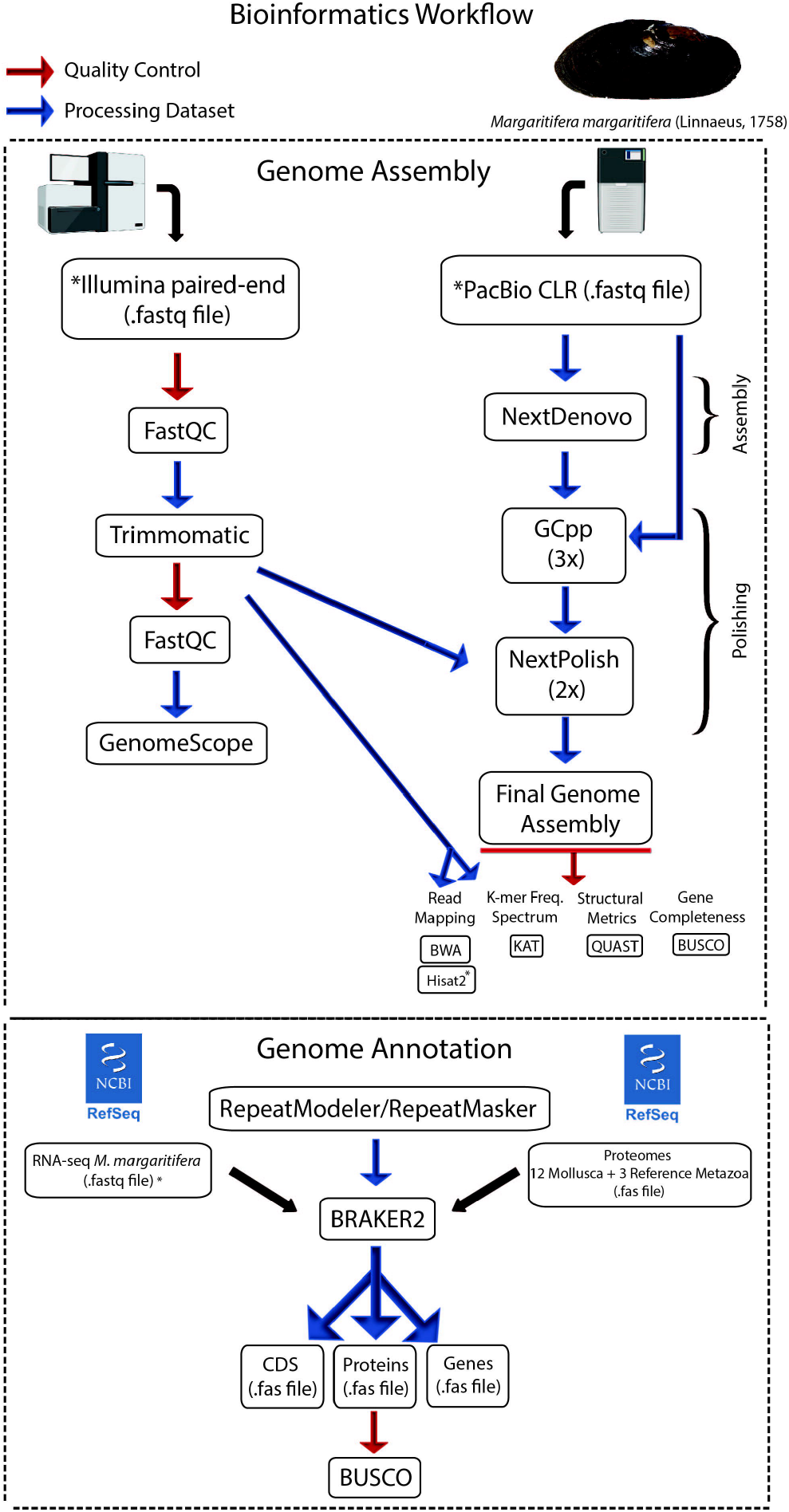
Bioinformatics pipeline applied for the genome assembly and annotation.

### Genome size and heterozygosity estimation

Before the assembly, the characteristics of the genome were accessed with a k-mer frequency spectrum using the paired-end reads. First, the quality of the reads was evaluated using FastQC (https://www.bioinformatics.babraham.ac.uk/projects/fastqc/) and the reads were after quality trimmed with Trimmomatic v.0.38 [28], specifying the parameters “LEADING: 5 TRAILING: 5 SLIDINGWINDOW: 5:20 MINLEN: 36”. The quality of the clean reads was validated in FastQC and after used for genome size estimation with Jellyfish v.2.2.10. and GenomeScope2 [29] specifying the k-mer length of 21.

### Genome assembly

The primary genome assembly was constructed using the raw PacBio reads with NextDenovo v2.4.0 (https://github.com/Nextomics/NextDenovo), with default parameters and specifying an estimated genome size of 2.4Gbp. Polishing of the resulting assembly was performed, first using PacBio reads, with three iterations of GCpp v 2.0.2 (Pacific Biosciences 2019), and after using the clean paired-end reads with two iterations of NextPolish v 1.2.3 [30]. PacBio read alignments were performed with pbmm2 v 1.4.0 (Pacific Biosciences 2019) and paired-end read alignments were performed with Burrows-Wheeler Aligner (BWA) v.0.7.17 [31], both with default parameters.

The general statistic and completeness of the final genome assembly were estimated with QUAST v5.0.2 [32], BUSCO v5.2.2 [33] and using the paired-end reads for read-back mapping, with BWA, and k-mer frequency distribution analysis with the K-mer Analysis Toolkit [34].

### Masking of repetitive elements, gene models predictions, and annotation

To mask repetitive elements, first, a *de novo* library of repeats was created for final genome assembly with RepeatModeler v.2.0.133 [35]. Subsequently, the genome was soft masked with RepeatMasker v.4.0.734 [36] combining the *de novo* library with the ‘Bivalvia’ libraries from Dfam_consensus-20170127 and RepBase-20181026.

Gene prediction was performed on the soft masked genome assembly using BRAKER2 pipeline v2.1.6 [37]. First, all the available RNA-seq data from *M. margaritifera* from GenBank [22,38] and Gomes-dos-Santos, et al., [18] (the same individual used for the genome assembly) was retrieved and quality trimmed with Trimmomatic v.0.38 (parameters described above). Afterwards, the clean reads were aligned to the masked genome, using Hisat2 v.2.2.0 with the default parameters [39]. Furthermore, the complete proteomes of 14 mollusc species and three reference Metazoa species (*Homo sapiens, Ciona intestinalis, Strongylocentrotus purpuratus*), downloaded from public databases (Table 3), were used as additional evidence for gene prediction. The BRAKER2 pipeline was then applied, specifying parameters “–etpmode; – softmasking;”. The gene predictions file (gff3) was renamed (Mma), cleaned, and filtered using AGAT v.0.8.0 [40], correcting overlapping prediction and removing incomplete gene predictions (i.e., without start and/or stop codons). Finally, proteins were extracted from the genome with AGAT and functional annotation was performed using InterProScan v.5.44.80 [41] and BLASTP searches against the RefSeq database [42]. Homology searches were performed using DIAMOND v.2.0.11.149 [43], specifying the parameters “-k 1, -b 20, -e 1e-5, - -sensitive, --outfmt 6”. Finally, BUSCO scores were estimated for the predicted proteins [33].

**Table 3.** List of proteomes used for BRAKER2 gene prediction pipeline.

| Phylum | Class | Order | Species | GenBank/RefSeq |
| --- | --- | --- | --- | --- |
| **Mollusca** | **Bivalves** |  |  |  |
|  |  | **Ostreida** |  |  |
|  |  |  | *Crassostrea gigas* | GCF_902806645.1 |
|  |  |  | *Crassostrea virginica* | GCF_002022765.2 |
|  |  | **Pectinida** |  |  |
|  |  |  | *Mizuhopecten yessoensis* | GCF_000457365.1 |
|  |  |  | *Pecten maximus* | GCF_902652985.1 |
|  |  | **Veneroida** |  |  |
|  |  |  | *Dreissena polymorpha* | GCF_020536995.1 |
|  |  |  | *Mercenaria mercenaria* | GCF_014805675.1 |
|  |  | **Unionida** |  |  |
|  |  |  | *Margaritifera margaritifera* | GCA_015947965.1 |
|  |  |  | *Megalonaias nervosa* | GCA_016617855.1 |
|  | **Gastropod** |  |  |  |
|  |  |  | *Biomphalaria glabrata* | GCF_000457365.1 |
|  |  |  | *Pomacea canaliculata* | GCF_003073045.1 |
|  |  |  | *Gigantopelta aegis* | GCF_016097555.1 |
|  | **Cephalopod** |  |  |  |
|  |  |  | *Octopus bimaculoides* | GCF_001194135.1 |
|  |  |  | *Octopus sinensis* | GCF_006345805.1 |
|  | **Polyplacophora** |  |  |  |
|  |  |  | *Acanthopleura granulata* | GCA_016165875.1 |
| **Chordata** |  |  | *Homo sapiens* | GCF_000001405.40 |
| **Chordata** |  |  | *Ciona intestinalis* | GCF_000224145.3 |
| **Echinodermata** |  |  | *Strongylocentrotus purpuratus* | GCF_000002235.4 |

### Data Validation

#### Sequencing results and genome assembly

The raw sequencing outputs resulted in a total of 103 Gbp of raw PacBio and 203 Gbp of raw paired-end reads. A total of 201 Gbp of paired-end reads were maintained after trimming and quality filtering. Similarly, to the results of [13], GenomeScope2 estimated genome size was ∼2.36 Gb and heterozygosity levels were low, i.e., ∼0.163 % (Figure 3a).

**Figure 3.**
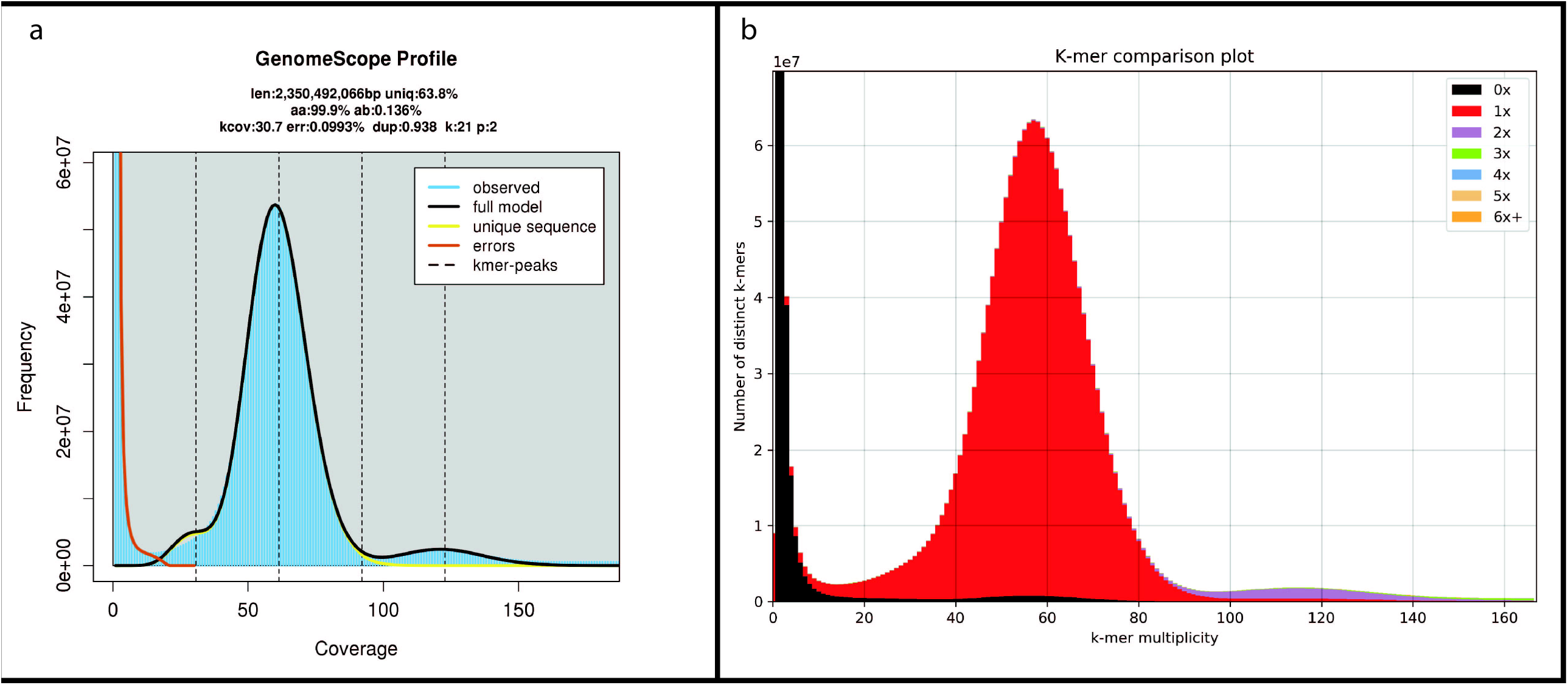
(a) GenomeScope2 k-mer (21) distribution displaying the estimation of genome size (len), homozygosity (aa), heterozygosity (ab), mean coverage of k-mer for heterozygous bases (kcov), read error rate (err), average rate of read duplications (dup), size of the k-mer used on the run (k:), and ploidy (p:). (b) *Margaritifera margaritifera* genome assembly assessment using KAT comp tool to compare the Illumina paired-end k-mer content within the genome assembly. Different colours represent the read k-mer frequency in the assembly.

The final genome assembly (hereafter referred to as Genome V2) has a total size of 2.45 Gbp, similar to the genome size reported in the previous assembly from [13] (hereafter referred to as Genome V1). Regarding the contiguity, Genome V2 shows a contig N50 of 3.42Mbp (Table 4), which represents a ∼202-fold increase in contig N50 and ∼11-fold increase in scaffold N50 relative to Genome V1 (Table 4). Additionally, Genome V2 represents the most contiguous freshwater mussel genome assembly currently available [13,25,44,45]. Genome V2 shows a ∼1.66-fold increase in N50 length regarding the other PacBio-based genome assembly, i.e., from *P. streckersoni*, [25], which is relevant considering that the Genome V2 is larger (nearly 4Mbp longer), has more repetitive elements (nearly 7% more) (Tables 4-5) and similar heterozygosity (nearly 0.43% less) (Figure 3).

**Table 4.**
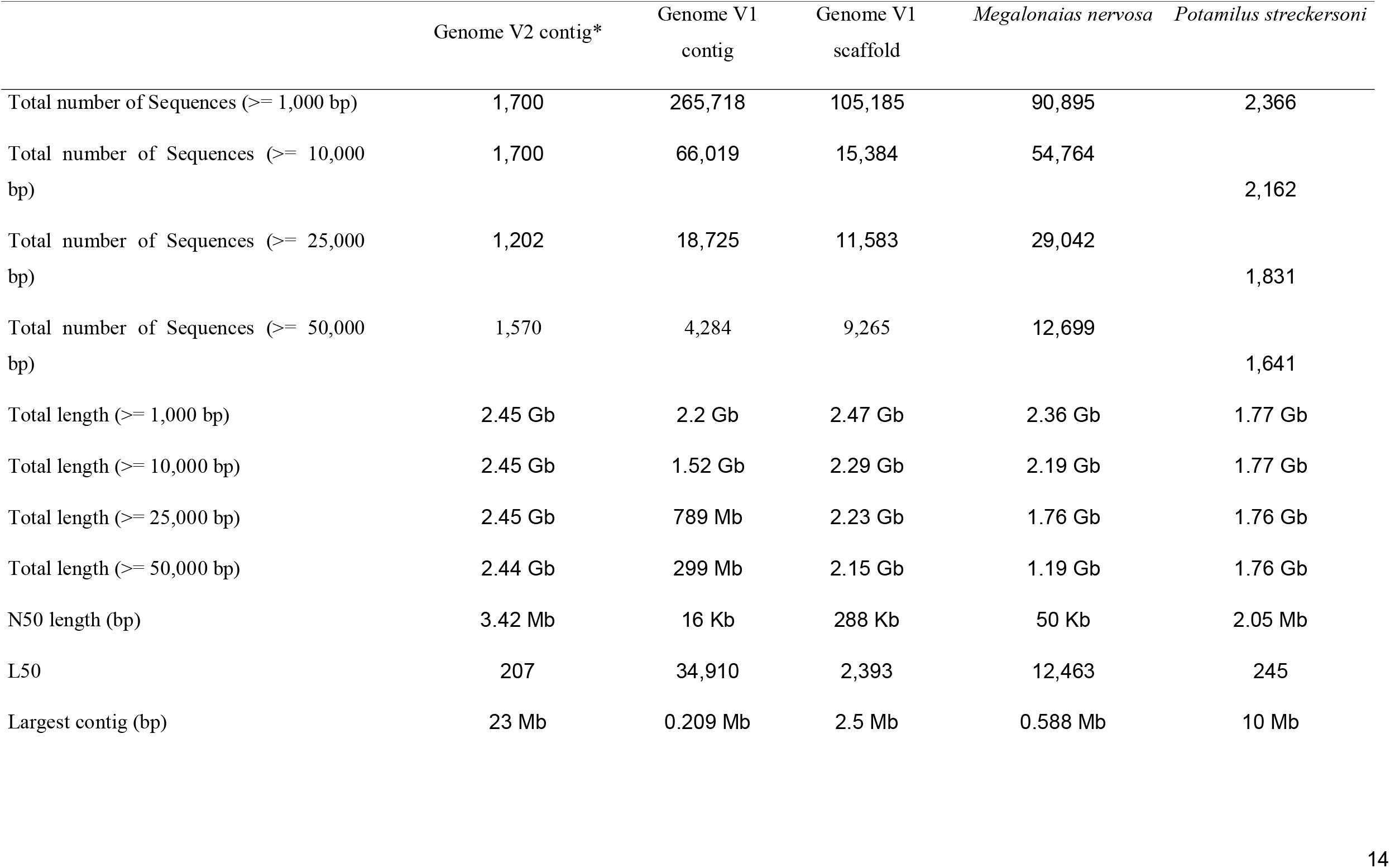
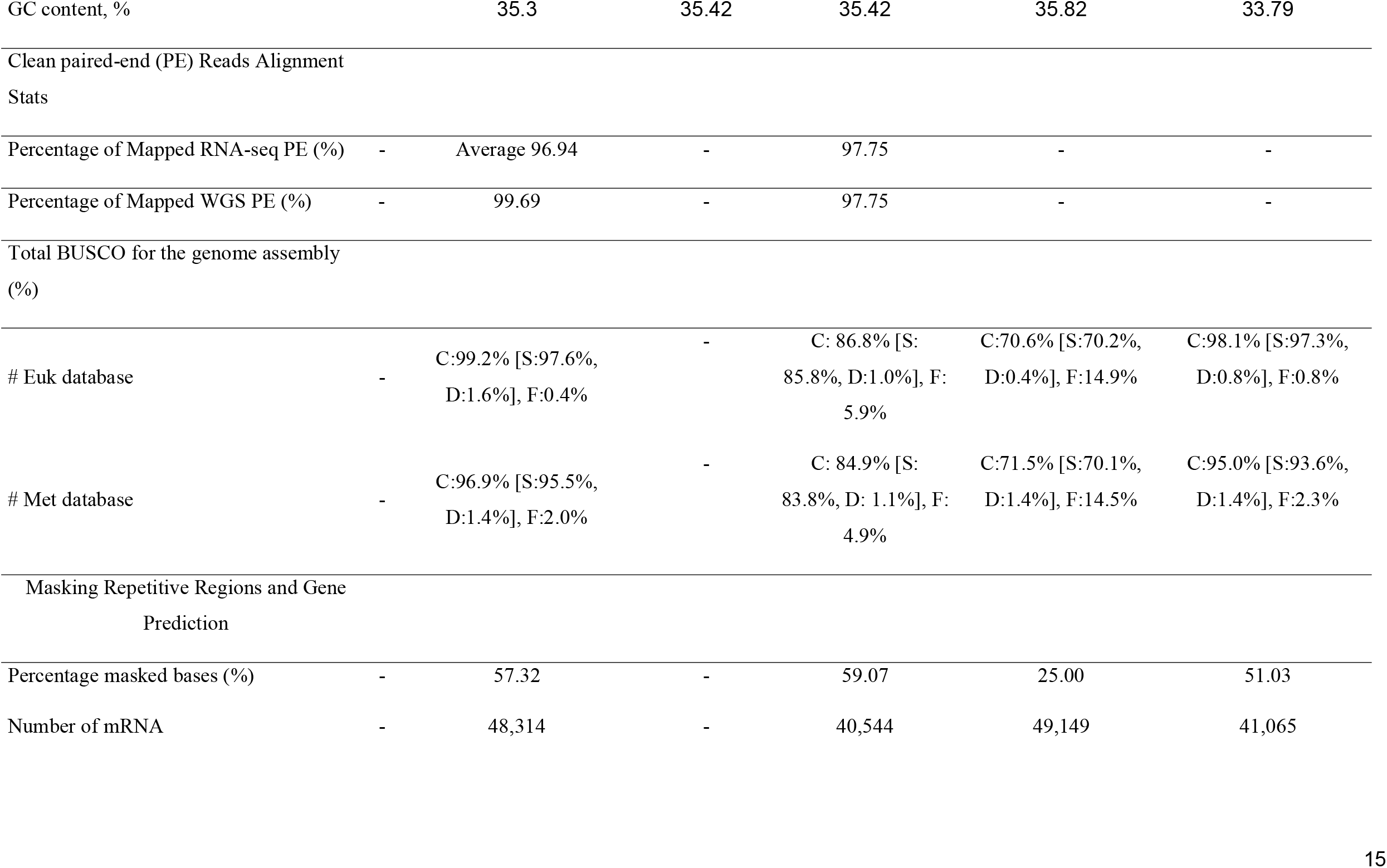
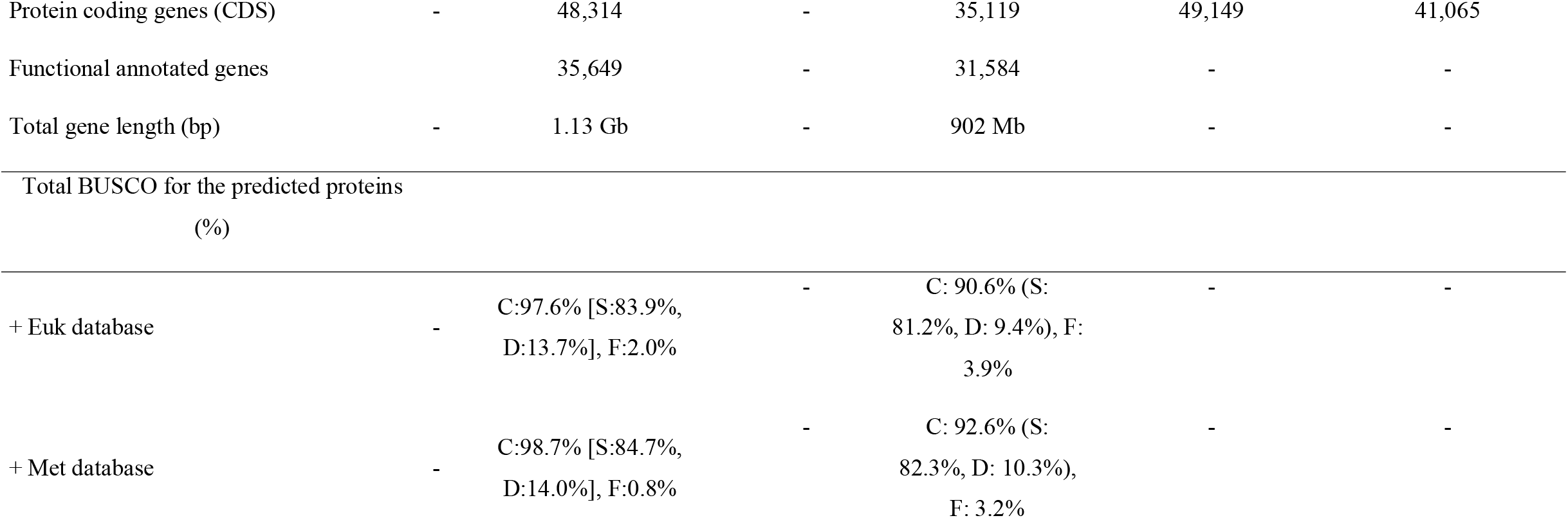
General statistics of the two *Margaritifera margaritifera* genome assemblies, including read alignment, gene prediction, and annotation. * Genome V2 refers to the new assembly here produced and is solely at the contig level, i.e., has no scaffolds; Genome V1 refers to the first *Margaritifera margaritifera* genome produced in [13] # Euk: From a total of 303 genes of Eukaryota library profile; # Met: From a total of 978 genes of Metazoa library profile; + Euk: From a total of 255 genes of Eukaryota library profile; + Met: From a total of 954 genes of Metazoa library profile; #,+ C: Complete; S: Single; D: Duplicated; F: Fragmented.

Genome V2 also shows a considerable increase in the BUSCOs scores, with nearly no fragmented nor missing hits for both the eukaryotic and metazoan curated lists of near-universal single-copy orthologous (Table 4). Short-read back-mapping percentages resulted in almost complete read mapping, 99.69% alignment rate (Table 4), and KAT k-mer distribution spectrum revealed that almost all read information was included in the final assembly (Figure 3b). Overall, these general statistics validate the high completeness, the low redundancy, and quality of the Genome V2.

### Repeat masking, gene models prediction, and annotation

RepeatModeler/RepeatMasker masked 57.32% of Genome V2 which is 1.75% less than the values of Genome V1, likely a consequence of the new assembly being able to resolve repetitive regions more accurately (Table 5). Furthermore, this value was considerably higher than the estimated duplications of GenomeScope, i.e., 36.2% (Figure 3a, Table 5). These differences have been observed in other assemblies of freshwater mussel genomes [4,25,44] and are likely a consequence of inaccurate estimation of repeat content when applying k-mer frequency spectrum analysis in highly repetitive genomes, using short reads. Similarly, to Genome V1, most repeats are unclassified (27.26%, ∼668Mgp), followed by DNA elements (17.18%, ∼421Mgp), long terminal repeats (5.95%, ∼145Mgp), long interspersed nuclear elements (5.86%, ∼143Mgp), and short interspersed nuclear elements (0.75%, ∼18Mgp) (Table 5). BRAKER2 gene prediction identified 48,314 CDS, which represents an increase compared with Genome V1, but is closer to the predictions of the other two freshwater mussel assemblies (Tables 4,6). This is probably a reflection of the higher contiguity and completeness of Genome V2, evidenced by high BUSCO scores for protein predictions, with almost no missing hits for either of the near-universal single-copy orthologous databases used (Table 3). The number of functionally annotated genes was also higher than those of Genome V1, with 4,065 additional genes annotated (Table 4,6,7). Overall, the numbers of both predicted and annotated genes are within the expected range for bivalves (reviewed in [4]), as well as within the records of other freshwater mussel assemblies [25,45].

**Table 5.**
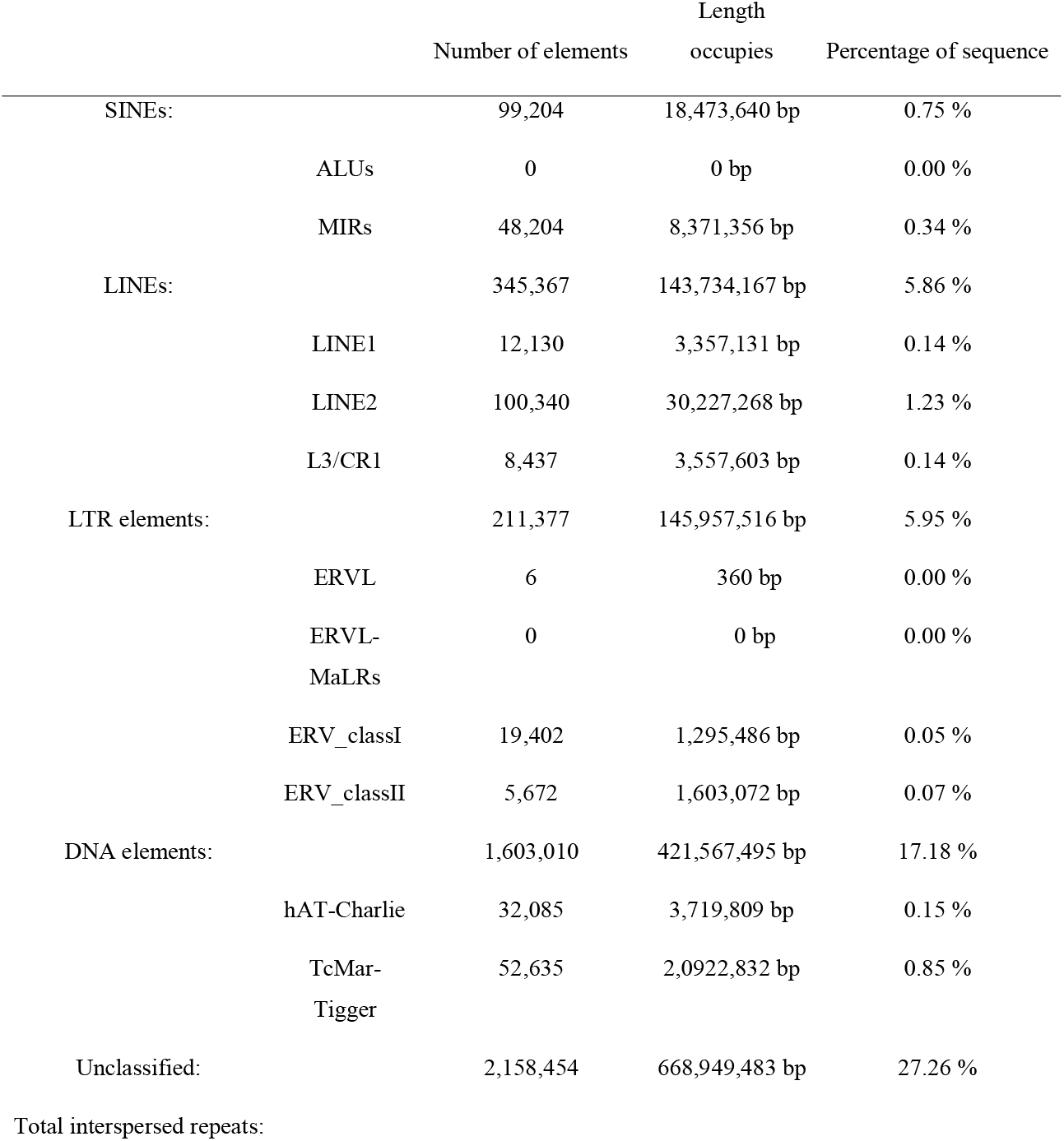
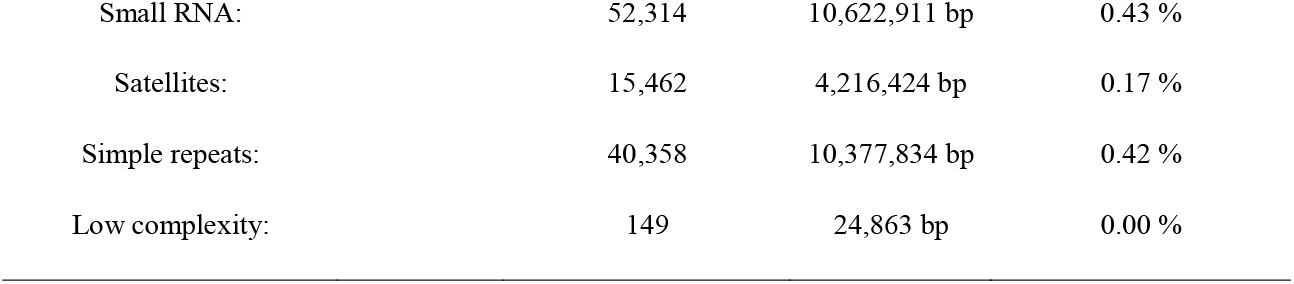
RepeatMasker report of the content of repetitive elements in the new *Margaritifera margaritifera* genome assembly.

|  |  | Number of elements | Length<br>occupies | Percentage of sequence |
| --- | --- | --- | --- | --- |
| SINEs: |  | 99,204 | 18,473,640 bp | 0.75 % |
|  | ALUs | 0 | 0 bp | 0.00 % |
|  | MIRs | 48,204 | 8,371,356 bp | 0.34 % |
| LINEs: |  | 345,367 | 143,734,167 bp | 5.86 % |
|  | LINE1 | 12,130 | 3,357,131 bp | 0.14 % |
|  | LINE2 | 100,340 | 30,227,268 bp | 1.23 % |
|  | L3/CR1 | 8,437 | 3,557,603 bp | 0.14 % |
| LTR elements: |  | 211,377 | 145,957,516 bp | 5.95 % |
|  | ERVL | 6 | 360 bp | 0.00 % |
|  | ERVL-MaLRs | 0 | 0 bp | 0.00 % |
|  | ERV_classI | 19,402 | 1,295,486 bp | 0.05 % |
|  | ERV_classII | 5,672 | 1,603,072 bp | 0.07 % |
| DNA elements: |  | 1,603,010 | 421,567,495 bp | 17.18 % |
|  | hAT-Charlie | 32,085 | 3,719,809 bp | 0.15 % |
|  | TcMar-Tigger | 52,635 | 2,0922,832 bp | 0.85 % |
| Unclassified: |  | 2,158,454 | 668,949,483 bp | 27.26 % |
| Total interspersed repeats: |  |  |  |  |

|  |  |  |  |
| --- | --- | --- | --- |
| Small RNA: | 52,314 | 10,622,911 bp | 0.43 % |
| Satellites: | 15,462 | 4,216,424 bp | 0.17 % |
| Simple repeats: | 40,358 | 10,377,834 bp | 0.42 % |
| Low complexity: | 149 | 24,863 bp | 0.00 % |

**Table 6.** Structural annotation report of the new *Margaritifera margaritifera* genome assembly.

| Structural Annotation | Number |
| --- | --- |
| Number of genes | 40,165 |
| Number of mrnas | 48,314 |
| Number of cdss | 48,314 |
| Number of exons | 328,489 |
| Number of introns | 280,173 |
| Number of start_codons | 48,314 |
| Number of stop_codons | 48,314 |
| Number of exon in cds | 328,489 |
| Number of intron in cds | 280,175 |
| Number of intron in exon | 280,175 |
| Number of intron in intron | 240,261 |
| Number gene overlapping | 440 |
| Number of single exon gene | 8,092 |
| Number of single exon mrna | 8,402 |
| mean mrnas per gene | 1.2 |
| mean cdss per mrna | 1.0 |
| mean exons per mrna | 6.8 |
| mean introns per mrna | 5.8 |
| mean exons per cds | 6.8 |
| mean introns in cdss per mrna | 5.8 |
| mean introns in exons per mrna | 5.8 |
| mean introns in introns per mrna | 5.0 |
| Total gene length | 1,134,996,674 |
| Total mrna length | 1,399,972,668 |
| Total cds length | 65,168,232 |
| Total exon length | 65,168,232 |
| Total intron length | 1,334,804,372 |
| Total start_codon length | 144,942 |
| Total stop_codon length | 144,942 |
| Total intron length per cds | 1,334,804,436 |
| Total intron length per exon | 1,334,804,436 |
| Total intron length per intron | 38,816,274 |
| mean gene length | 28,258 |
| mean mrna length | 28,976 |
| mean cds length | 1348 |
| mean exon length | 198 |
| mean intron length | 4,764 |
| mean cds piece length | 198 |
| mean intron in cds length | 4,764 |
| mean intron in exon length | 4,764 |
| mean intron in intron length | 161 |
| Longest gene | 492,278 |
| Longest mrna | 492,278 |
| Longest cds | 50,892 |
| Longest exon | 14,931 |
| Longest intron | 270,677 |
| Longest cds piece | 14,931 |
| Longest intron into cds part | 270,677 |
| Longest intron into exon part | 270,677 |
| Longest intron into intron part | 14,931 |
| Shortest gene | 123 |
| Shortest mrna | 123 |
| Shortest cds | 9 |
| Shortest exon | 3 |
| Shortest intron | 33 |

**Table 7.** Functional annotation report of *Margaritifera margaritifera* genome assembly.

| Functional Annotation | Number |
| --- | --- |
| Swissprot / RefSeq | 21,050 |
| CDD | 11,151 |
| Coils | 7,148 |
| GO | 18,024 |
| Gene3D | 23,613 |
| Hamap | 383 |
| InterPro | 29,030 |
| KEGG | 1,495 |
| MetaCyc | 1,445 |
| MobiDBLite | 12,555 |
| PIRSF | 973 |
| PRINTS | 5,250 |
| Pfam | 25,322 |
| ProSitePatterns | 6,471 |
| ProSiteProfiles | 15,115 |
| Reactome | 5,837 |
| SFLD | 110 |
| SMART | 12,494 |
| SUPERFAMILY | 23,283 |
| TIGRFAM | 1,148 |
| Total | 34,137 |

### Conclusion

In this report, a new and highly improved genome assembly for the freshwater pearl mussel is presented. This genome assembly, produced using PacBio long-read sequencing, represents a significant improvement in contiguity without requiring scaffolding. Unlike other freshwater mussels’ genomes, the one presented here has not been scaffolded (i.e., with no gaps of undetermined size), thus representing an ideal framework to employ chromosome anchoring approaches, such as Hi-C sequencing. This new genome represents a key resource to start exploring the many biological, ecological, and evolutionary features of this highly threatened group of organisms, for which the availability of genomic resources still falls far behind other molluscs.

### Data Records

Data has been uploaded to figshare (https://doi.org/10.6084/m9.figshare.22048250), which includes final unmasked and masked genome assemblies (Mma.fa and Mma_SM.fa), the annotation file (Mma_annotation_v1.gff3), predicted genes (Mma_genes_v1.fasta), predicted messenger RNA (Mma_mrna_v1.fasta), predicted open reading frames (Mma_cds_v1.fasta), predicted proteins (Mma_proteins_v1.fasta), as well as full table reports for Braker gene predictions and InterProScan functional annotations (Mma_annotation_v1_InterPro_report.txt) and RepeatMasker predictions (Mma_annotation_v1_RepeatMasker.tbl).

All software with respective versions and parameters used for producing the resources here presented (i.e., transcriptome assembly, pre and post-assembly processing stages, and transcriptome annotation) are listed in the methods section. Software programs with no parameters associated were used with the default settings.

## Declarations

## List of abbreviations

AGAT: Another Gtf/Gff Analysis Toolkit
BUSCO: Benchmarking Universal Single-Copy Ortholog
BWA: Burrows-Wheeler Aligner
CDS: Coding sequences
KAT: Kmer analyses toolkit
Gbp: gigabase pair(s)
Mbp: megabase pair(s)
kbp: kilobase pair(s)
mtDNA: NCBI: National Center for Biotechnology Information
NT-NCBI: Nucleotide database of NCBI
PacBio: Pacific Biosciences
PE: paired-end
QUAST: Quality Assessment Tool for Genome Assemblies
SMRT: Single Molecule, Real-Time

## Ethical approval

This work has been approved by the CIIMAR ethical committee and by CIIMAR Managing Animal Welfare Body (ORBEA) according to the European Union Directive 2010/63/EU.

## Author contributions

E.F, M.L.L, L.F.C.C designed and conceived this work.

M.L.L, VP and A.T were responsible for the field sampling.

A.G.S and A.M.M carried out the bioinformatics analyses.

L.F.C.C, EF, GA and TF revised the bioinformatics analyses.

A. G. S. and E. F wrote the first version of the manuscript.

All authors read, revised, and approved the final manuscript.

## Acknowledgments

Not applicable.

